# Crystal structure and functional peculiarities of a primordial Orange Carotenoid Protein (OCPX)

**DOI:** 10.1101/2022.06.14.496144

**Authors:** Yury B. Slonimskiy, Andrey O. Zupnik, Larisa A. Varfolomeeva, Konstantin M. Boyko, Eugene G. Maksimov, Nikolai N. Sluchanko

**Affiliations:** A.N. Bach Institute of Biochemistry, Federal Research Center of Biotechnology of the Russian Academy of Sciences, 119071 Moscow, Russian Federation; M.V. Lomonosov Moscow State University, Faculty of Biology, 119991 Moscow, Russian Federation

**Keywords:** Orange Carotenoid Protein X, oligomeric state, photoactivity, structure, SEC-MALS

## Abstract

The two-domain photoactive Orange Carotenoid Protein (OCP) confers photoprotection in cyanobacteria and presumably stems from domain fusion. Yet, the primitive thylakoid-less cyanobacteria *Gloeobacter* encodes a complete OCP. Its photosynthesis regulation lacks the so-called Fluorescence Recovery Protein (FRP), which in *Synechocystis* inhibits OCP-mediated phycobilisome fluorescence quenching, and *Gloeobacter* OCP belongs to the recently defined, heterogeneous clade OCPX (GlOCPX), the least characterized compared to OCP2 and especially OCP1 clades. Here we describe the first crystal structure of OCPX and provide its detailed structural and functional comparison with OCP1 from *Synechocystis*. Monomeric GlOCPX quenches *Synechocystis* phycobilisomes but displays drastically accelerated, less temperature-dependent recovery after photoactivation, evades regulation by FRP from other species and reveals numerous structural features reflecting its functional peculiarities. Our detailed description of a primordial OCPX sheds light on the evolution of the OCP-dependent photoprotection mechanism, rationalizing subdivision of the OCPX clade into subclades.

## Introduction

High light conditions require the presence in photosynthetic organisms of regulatory mechanisms protecting the cell against the photodamage and reactive oxygen species (ROS) that can form upon absorption of excess light by light-harvesting complexes ^1^. Glaucophytes, red algae and most cyanobacteria have unique soluble antenna complexes called the phycobilisomes (PBs) ^2^, megadalton phycobiliprotein assemblies that attach to the photosystems and transfer energy to their reaction centers, improving light harvesting efficiency ^3,4^. PBs-containing cyanobacteria have developed a unique photoprotection system involving a water-soluble ∼35 kDa photoactive Orange Carotenoid Protein (OCP) that serves simultaneously a blue-light sensor and effector of non-photochemical quenching (NPQ) ^5–7^.

OCP is a unique photoreceptor whose photoactivity is determined by the carotenoid cofactor, a ketocarotenoid 3’-hydroxyechinenone (hECH), echinenone (ECH) or canthaxanthin (CAN) ^5,8,9^. Although OCP can bind hydroxycarotenoids, only OCP complexes with ketocarotenoids are photoactive ^9,10^. According to X-ray structures of OCP known since 2003 ^11^, such complexes are formed upon embedment of the ketocarotenoid between two structural domains, the N-terminal (NTD) and C-terminal (CTD) connected by a long interdomain linker ^12–14^. The β-ionone rings of the carotenoid are placed in both domains, yet only in the CTD the keto-group of the carotenoid is coordinated by H-bonds with Trp288 and Tyr201 side chains (*Synechocystis* OCP numbering) ^12–14^. The opposite β-ionone ring of the carotenoid may lack a ketogroup and contacts the Trp110 and Tyr44 side chains (*Synechocystis* OCP numbering) ^13,14^. The compact two-domain structure of OCP is stabilized by the N-terminal extension (NTE) attached to the CTD, which, together with the specific salt-bridge connecting the NTD and CTD (R155-E244 in SynOCP), plays an important role in OCP photoactivation ^14–18^.

Photon absorption by dark-adapted, orange OCP state (OCP^O^.) entails a chain of photochemical and protein transformations involving i) breaking of H-bonds between the carotenoid and key residues in the CTD, ii) sliding of the carotenoid from its original position completely into the NTD, iii) NTE detachment from its binding site on the CTD surface and iv) a complete domain separation ^15,19–23^. The photoactivated, red OCP state (OCP^R^) has an expanded size due to the separated NTD and CTD ^21,24–26^; it binds to the PBs core, which results in quenching of their fluorescence by dissipation of the excess absorbed energy into heat ^5,27^. The low quantum yield of the photoconversion ensures that only high light conditions activate NPQ. According to time-resolved studies, OCP photoactivation involves numerous intermediates having distinct spectral and structural properties (see i-iv above) ^18,19^. The light-triggered accumulation of the PBs-quenching OCP^R^ form is spontaneously reversed in the dark, which is a highly temperature-dependent process ^5,13,18,19,24^.

In *Synechocystis, Anabaena* and *Arthrospira* and many other cyanobacteria, OCP^R^-OCP^O^. transition is drastically accelerated by the so-called Fluorescence Recovery Protein (FRP) ^28,29^. FRP is a ∼13 kDa dimeric α-helical protein ^30^ that specifically binds to the CTD of the photoactivated OCP and promotes OCP relaxation ^29–31^. FRP therefore inhibits the OCP-mediated NPQ, which is no longer required in low light, when efficient light harvesting is again a priority ^28,30–32^. Interestingly, FRP homologs are also found in non-cyanobacteria and even in non-photosynthetic prokaryotes, suggesting that they have additional functions beyond photoprotection and that their role in photoprotection is rather young ^33^. While low-homology FRPs from different cyanobacteria can act on *Synechocystis* OCP, their interaction with OCP can differ by efficiency and prevalent stoichiometry ^34^.

The canonical photoprotection system of cyanobacteria composed of PBs, OCP and FRP can be reconstituted *in vitro* ^31^. Nevertheless, its functioning has mainly been studied only for a handful of model organisms (e.g., *Synechocystis, Anabaena* and *Arthrospira*). Recent bioinformatic works revealed that many cyanobacteria may lack FRP, may have independently encoded functional homologs of the NTD (HCP, helical carotenoid proteins, clades 1-9 ^35^) and CTD domains of OCP (CTDH, C-terminal domain homologs, clades 1 and 2 ^36^), and that the OCP family contains at least three different clades ^37–39^. A given cyanobacterium can have a different set of above-mentioned genes (including a mixture of various OCP types), or can lack one of the proteins at all. The evolutionary aspects and functions of individual proteins related to the OCP-dependent photoprotection system remain controversial. The existence of NTD and CTD homologs as individual carotenoproteins suggested that the two-domain OCP could arise in the evolution as the result of domain fusion ^37,38,40,41^. Recent analysis of OCP diversity and phylogenetics revealed that among the three currently considered OCP clades (OCP1, OCP2 and OCPX ^37^), the OCP1 and OCP2 branches diverged only long after their common ancestor split off from the most ancient OCPX clade ^38^. OCP1, OCP2 and OCPX are thought to differ by the rate of OCP^R^-OCP^O^. recovery, oligomeric status and the ability to be regulated by FRP ^37,38^. Notwithstanding, the representatives of OCP clades are disproportionately studied. For example, there is only one publication on OCPX characterization ^38^. In contrast to at least six non-redundant crystal structures of OCP1 from different cyanobacteria available (Table 1), structural data on OCP2 and OCPX representatives were not reported as these proteins proved difficult to crystallize ^42^.

**Table 1.** Non-redundant crystal structures of OCP available to date.

| PDB ID | Clade | Organism | Carotenoid | Resolution | Crystallographic assembly unit | Ref. |
| --- | --- | --- | --- | --- | --- | --- |
| 6PQ1 | OCP1 | *Fremyella diplosiphon* (Tolypothrix sp. PCC 7601) | CAN | 1.61 Å | dimer | <sup>42</sup> |
| 5HGR | OCP1 | *Anabaena* (Nostoc) sp. PCC 7120 | CAN | 1.68 Å | dimer | <sup>69</sup> |
| 7EKR | OCP1 | *Gloeocapsa* sp. PCC 7513 | ECH | 2.17 Å | dimer | n/a |
| 5UI2 | OCP1 | *Limnospira* (Arthrospira) maxima | hECH | 2.10 Å | dimer | <sup>11</sup> |
| 3MG1 | OCP1 | *Synechocystis* sp. PCC 6803* | ECH | 1.65 Å | dimer | <sup>14</sup> |
| 7QD2 | OCP1 | *Planktothrix* PCC7805** | CAN | 1.40 Å | dimer | <sup>70</sup> |
| 8A0H | OCPX | *Gloeobacter kilaueensis* JS1 | ECH | 1.73 Å | monomer | this work |

\*Other non-mutant SynOCP1 structures are: 4XB5 (with CAN) <sup>13</sup>, 5TUW and 5TV0 (with hECH), and 5TUX (ECH) <sup>71</sup>.
\*\*The highest-resolution structure from a set including other structures of this OCP: 7QD0 (ECH, 1.70 Å), 7QCZ (CAN, 1.85 Å), 7QD1 (ECH, 1.71 Å) <sup>70</sup>.

Focusing on the primordial cyanobacterium *Gloeobacter kilaueensis* JS1, whose genome was sequenced in 2013 ^43^ and contains a single OCPX gene (according to the earlier classification ^37^), here we report the first crystal structure of OCPX and provide its detailed structural and functional comparison with OCP1 from *Synechocystis* sp. PCC 6803 (SynOCP1) (50.5% identity, see Fig. 1A). Our structural data explain the distinctive character of the most divergent OCPX group from *Gloeobacteria* and give new insights into the evolution of the OCP-related photoprotection mechanism of cyanobacteria, rationalizing further subdivision of the OCPX clade into subclades.

**Fig. 1.**
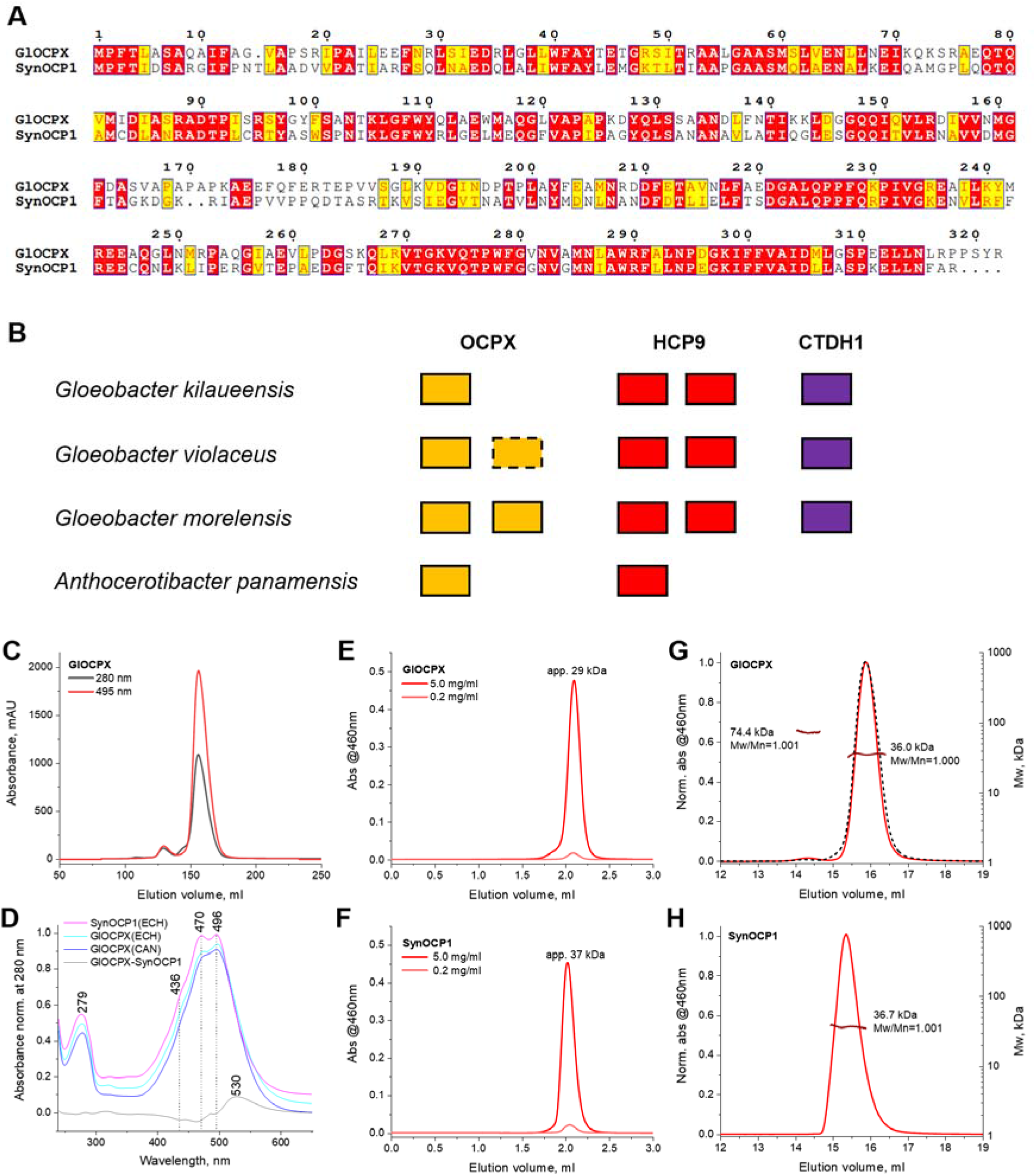
Comparison of GlOCPX and SynOCP1. A. Pairwise sequence alignment of GlOCPX (Uniprot U5QHX0) and SynOCP1 (Uniprot P74102). Identical residues are highlighted by red, similar - by yellow. B. Occurrence and number of OCP-related genes in the sequenced *Gloeobacteria* species. Clades classification complies with ^35,37^. A dashed line denotes an incomplete OCPX sequence lacking first ∼50 residues, which likely corresponds to a nonfunctional OCP protein. C. Preparative SEC of GlOCPX(CAN) on a Superdex 75 26/600 column showing two oligomeric forms. D. Absorbance spectra of GlOCPX with either ECH or CAN, compared with SynOCP1(ECH). The spectra correspond to the peak maxima on the SEC profiles and are shifted along the Y axis for clarity. A difference spectrum GlOCPX(ECH) minus SynOCP1(ECH) is also shown. E,F. Analytical SEC profiles of GlOCPX (E) and SynOCP1 (F) obtained on a Superdex 200 Increase 5/150 column using 25 times different protein concentrations in the loaded samples (indicated in mg/ml). Apparent Mw values corresponding to the peak maxima obtained using column calibration are indicated. G,H. SEC-MALS of GlOCPX (G) or SynOCP1 (H) (5 mg/ml, 50 μl) on a Superdex 200 Increase 10/300 column showing Mw distributions across the peaks along with the Mw/Mn polydispersity indices. Average Mw values for the distributions are indicated. A dashed line in panel G shows the profile for the highly concentrated GlOCPX sample (17 mg/ml, 50 μl).

## Results

### GlOCPX is arguably the most ancient extant OCP

*Gloeobacteria* is thought to be the most primitive extant cyanobacteria preserving many ancestral characteristics, including the lack of thylakoids and circadian clock, low capacity for maintaining a constant internal pH, bundle-shaped PBs and bacterial-type carotenoid biosynthesis ^44–47^. *Gloeobacteria* presumably diverged from the other cyanobacteria over 2 Ga years ago and are considered the basal cyanobacteria group ^43,48–50^. Among 4430 potential protein encoding genes, *Gloeobacter violaceus* has only 610 genes matching to other cyanobacterial genomes, whereas as many as 684 genes are unique to this ancient cyanobacterium ^51^. All four sequenced *Gloeobacteria* species already have at least one complete OCP gene (OCPX only), a couple HCP genes, a CTDH gene and no FRP (Fig. 1B). Such a primordial OCPX has never been characterized. At least one of the strains, *Gloeobacter violaceus* PCC 7421, was documented to express an OCPX (verified by immunoblotting and mass-spectrometry) and display blue-light-induced NPQ ^49^. All sequenced *Gloeobacter* species have genetic chassis for biosynthesizing ketocarotenoids (https://www.genome.jp/pathway/glj00906) required for OCP photoactivity. Moreover, *G. violaceus* PCC 7421 was reported to synthesize appreciable amounts of ECH by using its CrtW ketolase, in addition to other two of its endogenous carotenoids, β-carotene and oscillol difucoside ^45^. Together, these facts indicate that *Gloeobacter* OCPX (GlOCPX) is a functional photoprotective protein *in vivo*.

### GlOCPX forms stable monomer binding ketocarotenoid

Despite the attractiveness of *Gloeobacter* species for many physiological and biochemical studies, their use is limited by extremely low growth rates ^47^. Therefore, we produced GlOCPX in *Escherichia coli* strains synthesizing ketocarotenoids canthaxanthin (CAN) or echinenone (ECH) ^25,52^. On preparative SEC, GlOCPX yielded two oligomeric forms, a >20 times more abundant monomer and less abundant dimer (Fig. 1C), similar for ECH and CAN. Use of diode-array detection revealed that, compared to the SynOCP1 spectrum having nearly equal amplitude of its two maxima at 470 and 496 nm, the maximum at 496 nm was considerably higher for both GlOCPX with ECH and CAN, and the vibronic structure of the latter was least pronounced (Fig. 1D). The GlOCPX spectra were significantly red-shifted compared to SynOCP1. This agrees with ^38^ and may indicate differences in the steady-state carotenoid conformation. Of note, the absorbance spectrum of GlOCPX remains unchanged by the addition of kosmotropes (0.75 M phosphate, see Supplementary Fig. 1), which indicates that the red-shift is not caused by spontaneous domain separation and rather reflects the conformational mobility of the carotenoid β-ring in the CTD.

In contrast to the previous report that OCPX proteins are dimers ^38^, we could not detect any significant dimerization of the isolated GlOCPX monomer even upon its drastic concentration, which was the same for SynOCP1 (Fig. 1E,F). While the protein peak did not shift upon increasing protein concentration up to 17 mg/ml (Fig. 1G), the apparent Mw for GlOCPX (29 kDa) was appreciably smaller than that of SynOCP1 (37 kDa) and than that calculated from GlOCPX sequence (35.8 kDa). This uncertainty prompted us to use SEC-MALS to determine absolute Mw. Despite an obviously different position on the elution profile, GlOCPX peak was characterized by absolute Mw typical for the monomer (36 kDa), with excellent monodispersity (Fig. 1G). Absolute Mw for the similarly monodisperse SynOCP1 was 36.7 kDa (Mw calculated from sequence 34.6 kDa, Fig. 1H).

### Interface mutations affect GlOCPX dimerization and crystallization

More detailed analysis revealed a negligibly small fraction of GlOCPX dimer (absolute Mw 74.4 kDa; calculated dimeric Mw - 71.6 kDa) appeared after concentrating the monomer sample (Fig. 1G). Interestingly, all so far crystallized OCP belong to the OCP1 clade and formed crystallographic dimers (Table 1). Using the crystallographic SynOCP1 dimer as a template, we hypothesized that GlOCPX can form a similar interface and predicted its structure using Alphafold2 ^53^ (Supplementary Fig. 2A,B). This revealed that while the antiparallel SynOCP1 dimer features two interface alanines (Ala26), the equivalent positions in GlOCPX are occupied by Glu25 residues, which would lead to a severe clash disfavoring such interface. Consequently, an extensive crystal screening failed for GlOCPX WT, but turned successful for its E25A mutant, which tentatively imitated the SynOCP1 interface (Supplementary Fig. 2C).

Motivated by this, we prepared another, E25C mutant of GlOCPX that could be used for disulfide trapping of the tentative dimer interface under non-reducing conditions. In such dimer, the E25C mutation would bring Cβ atoms of cysteines to a 3.7 Å distance, sufficient for the formation of the disulfide bridge. Of note, GlOCPX does not have endogenous cysteines, therefore Cys25 would provide for an unambiguous interpretation. The GlOCPX-E25C mutant displayed a significantly increased dimer/monomer ratio (∼25%) (Supplementary Fig. 2D) compared to the WT protein (∼5%) (Fig. 1C). The two oligomeric forms revealed indistinguishable absorbance spectra (Supplementary Fig. 2E) and a perfect peak separation, suggesting that dimerization does not affect carotenoid microenvironment and that the corresponding dimer/monomer equilibrium is kinetically locked. Unexpectedly however, disulfide trapping of either the dimeric or monomeric E25C fraction produced only ∼10% of the cross-linked dimers (Supplementary Fig. 2F). Hence the expected dimeric interface is likely not populated by GlOCPX. To avoid heterogeneity associated with the cysteine presence, we switched back to the E25A mutant.

### Crystal structure of GlOCPX

While crystal screening for the WT protein and the E25A dimer was unsuccessful, the E25A monomer fraction produced crystals (Supplementary Fig. 2C) that diffracted to 1.7 Å, enabling structure solution (Table 2). Surprisingly, the asymmetric unit contained a GlOCPX monomer forming a lattice where the expected dimer assembly was not found (Fig. 2A). An apparent crystallization-promoting effect of the E25A mutation was explained by the location of Ala25 residues in crystal contacts that could not accommodate bulkier glutamate side chains (Fig. 2B). The orientation and contacts of neighboring GlOCPX monomers differed significantly from that in the crystallographic SynOCP1 dimer (Fig. 2C). The small area of new interfaces (<<900 Å^2^) suggested lack of biological relevance.

**Table 2.**
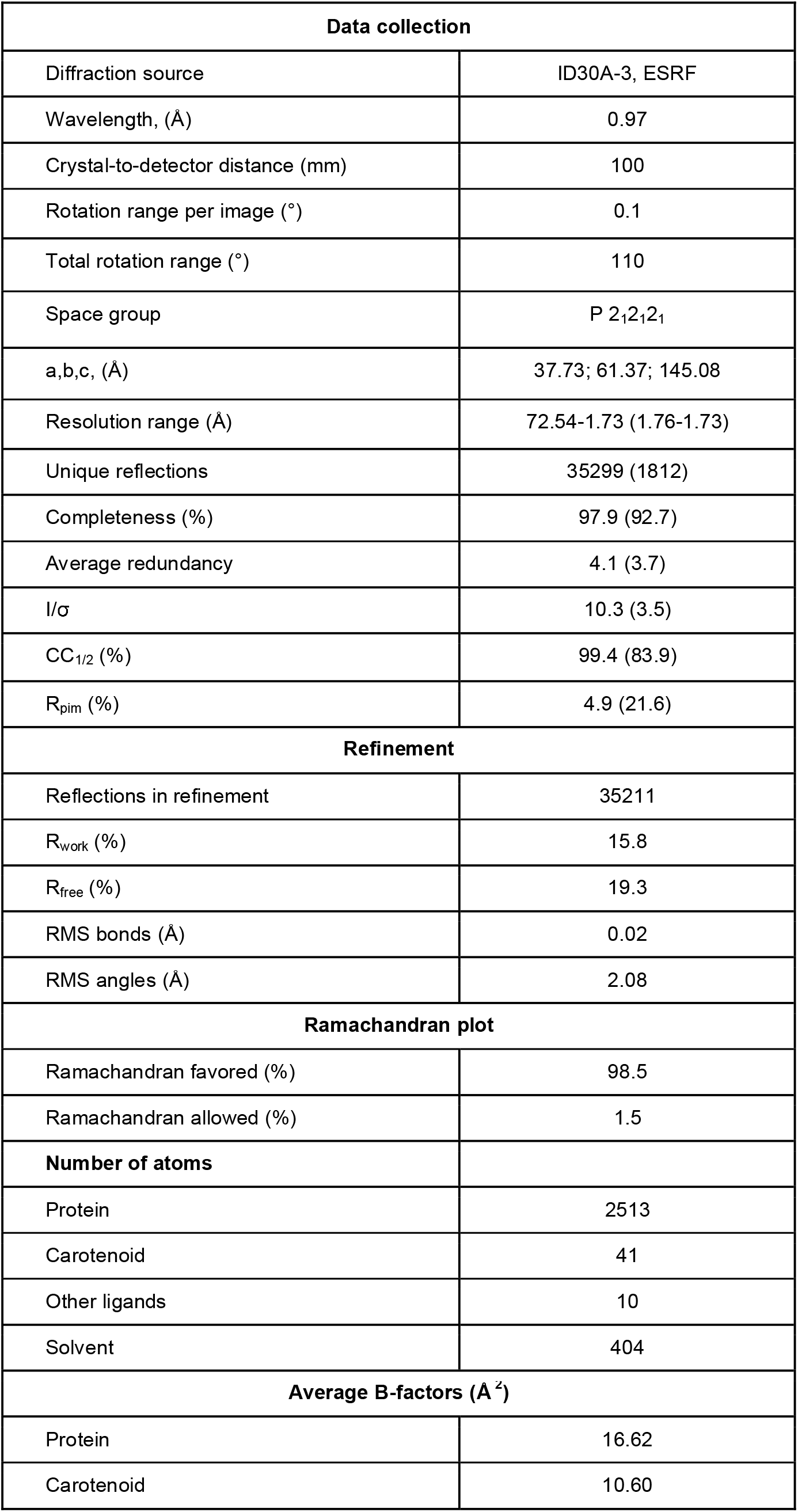
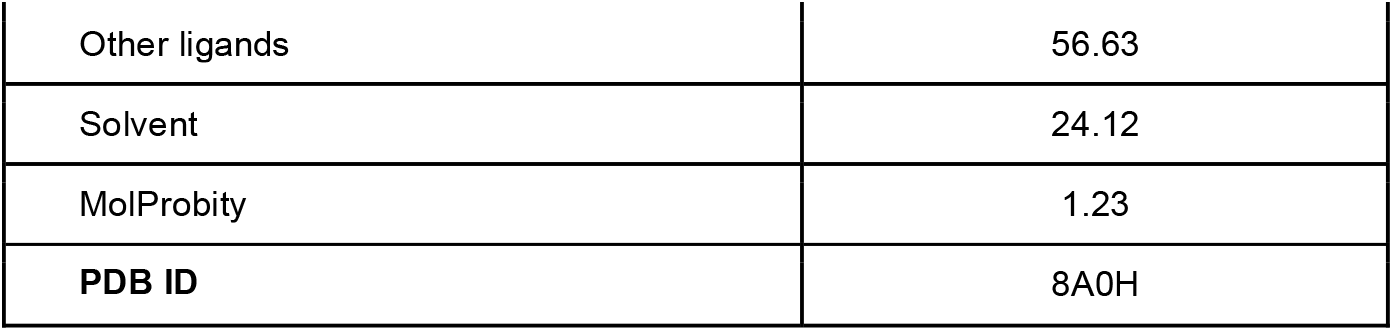
Diffraction data collection and refinement statistics for GOCPx E25A mutant.

| Data collection |  |
| --- | --- |
| Diffraction source | ID30A-3, ESRF |
| Wavelength, (Å) | 0.97 |
| Crystal-to-detector distance (mm) | 100 |
| Rotation range per image (°) | 0.1 |
| Total rotation range (°) | 110 |
| Space group | P 2<sub>1</sub>2<sub>1</sub>2<sub>1</sub> |
| a,b,c, (Å) | 37.73; 61.37; 145.08 |
| Resolution range (Å) | 72.54-1.73 (1.76-1.73) |
| Unique reflections | 35299 (1812) |
| Completeness (%) | 97.9 (92.7) |
| Average redundancy | 4.1 (3.7) |
| I/σ | 10.3 (3.5) |
| CC<sub>1/2</sub> (%) | 99.4 (83.9) |
| R<sub>pim</sub> (%) | 4.9 (21.6) |
| Refinement |  |
| Reflections in refinement | 35211 |
| R<sub>work</sub> (%) | 15.8 |
| R<sub>free</sub> (%) | 19.3 |
| RMS bonds (Å) | 0.02 |
| RMS angles (Å) | 2.08 |
| Ramachandran plot |  |
| Ramachandran favored (%) | 98.5 |
| Ramachandran allowed (%) | 1.5 |
| Number of atoms |  |
| Protein | 2513 |
| Carotenoid | 41 |
| Other ligands | 10 |
| Solvent | 404 |
| Average B-factors (Å<sup>2</sup>) |  |
| Protein | 16.62 |
| Carotenoid | 10.60 |
| Other ligands | 56.63 |
| Solvent | 24.12 |
| MolProbity | 1.23 |
| **PDB ID** | 8A0H |

**Fig. 2.**
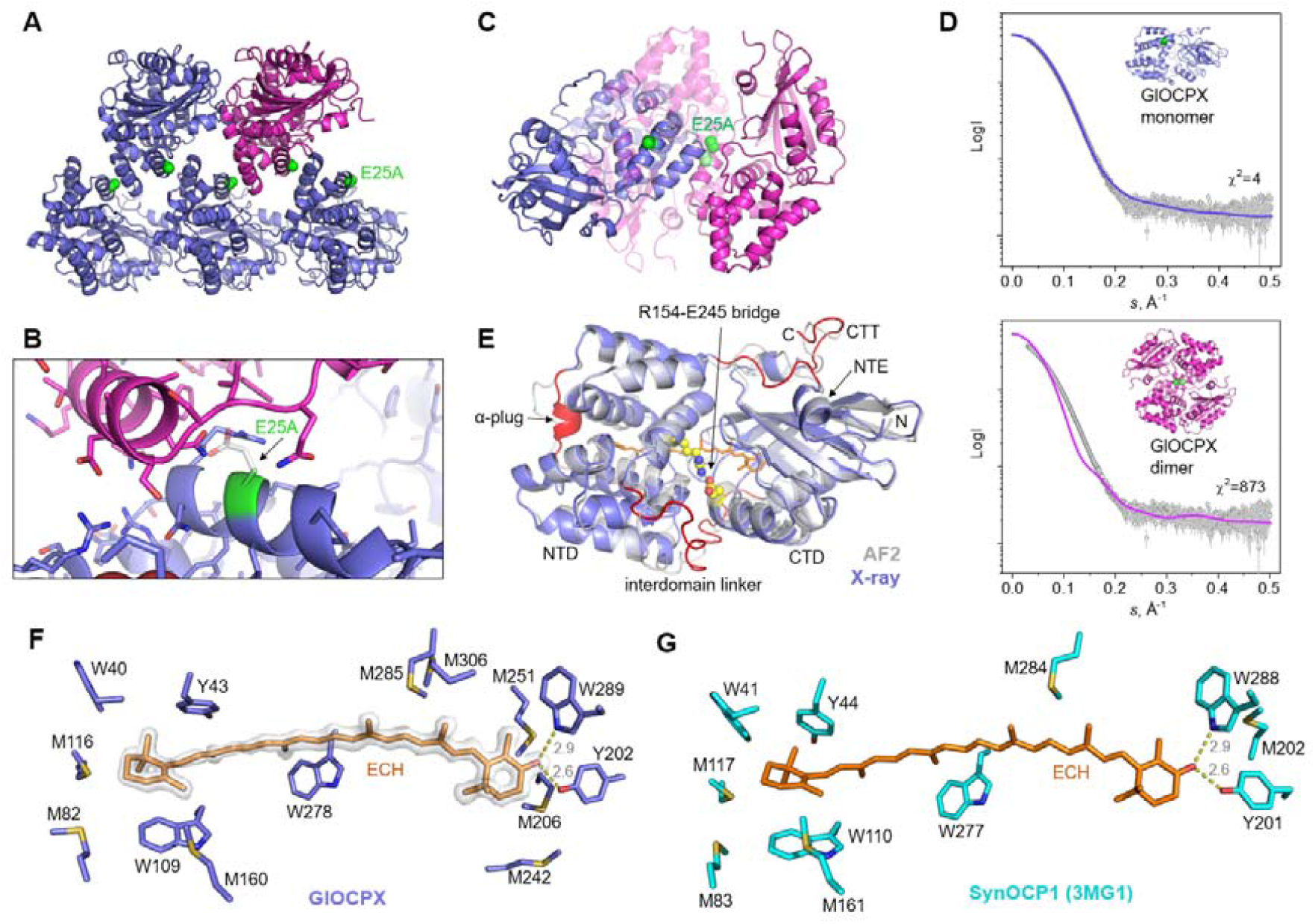
Crystal structure of the GlOCPX-E25A mutant. A. Crystal lattice showing the spatial arrangement of the mutated Ala25 residues (green spheres). One protein monomer is shown by magenta for clarity. B. A close-up view illustrating that the E25A mutation promotes crystal contact formation. Note that the bulky Glu25 residue (white sticks) could not be accommodated within such contacts. C. GlOCPX-E25A monomer orientations like those in SynOCP1 crystals (magenta cartoon) and in our crystal structure (magenta and violet cartoon). Ala25 residues are shown by green spheres. D. SAXS data for GlOCPX-E25E and their approximation by either GlOCPX monomer or SynOCP1-like crystallographic dimer. E. Superposition of GlOCPX-E25A crystal structure (violet cartoon) and its prediction by Alphafold2 ^53^ (semitransparent grey cartoon) showing novel features revealed by the crystal structure (red). The key interdomain salt bridge (yellow spheres) stabilizing the compact form of SynOCP1 is also conserved in GlOCPX. F,G. Echinenone microenvironment in GlOCPX (F) or in SynOCP1 (G). In F, the 2Fo-Fc electron density map fragment contoured at 1σ (white semitransparent surface) is also shown. Distances are shown in A.

To characterize the GlOCPX structure in solution, we applied small-angle X-ray scattering (SAXS). This unequivocally revealed that only the GlOCPX E25A monomer, and not the dimer, provides satisfactory description of the SAXS data (Fig. 2D). The monomeric GlOCPX state was evident from SAXS-derived structural parameters (*R*_g_=2.25 nm, *R*_max_=7.7 nm; Mw 32-35 kDa, Table 3).

**Table 3.** SAXS-derived structural parameters of GlOCPX.

| Parameter | GIOCPX WT | GIOCPX-E25A |
| --- | --- | --- |
| Mw calculated from sequence | 35.8 kDa | 35.8 kDa |
| Number of residues | 325 | 325 |
| Protein concentration | 17.5 mg/ml 35 μl loaded on a Superdex 200 Increase 3.2/300 column | 13.5 mg/ml 40 μl loaded on a Superdex 200 Increase 3.2/300 column |
| *R*<sub>g</sub> (Guinier) | 22.4 ± 0.1 Å * | 22.5 ± 0.2 Å |
| s*R*<sub>g</sub> limits | 0.67-1.29 | 0.73-1.29 |
| *R*<sub>g</sub> (reverse) | 22.5 ± 0.2 Å | 22.5 ± 0.1 Å |
| *D*<sub>max</sub> | 77 Å | 77 Å |
| Porod volume | 57 800 Å<sup>3</sup> | 56 800 Å<sup>3</sup> |
| *M*<sub>w</sub> Porod | 36.1 kDa | 35.5 kDa |
| *M*<sub>w</sub> MoW | 33.9 kDa | 36.6 kDa |
| *M*<sub>w</sub> Vc | 32.4 kDa | 32.8 kDa |
| Kratky plot | bell-shaped (folded) | bell-shaped (folded) |

\*All SAXS-derived parameters were calculated using the ATSAS 2.8 software package <sup>72</sup>.

While the crystal structure of GlOCPX was solved using an Alphafold2 model, the final refined model revealed multiple unpredicted features (Fig. 2E). These included a different conformation of the loop connecting the NTE with NTD, of the C-terminal tail (CTT), and the interdomain linker, which all are essential to the OCP function ^15,16,54,55^. The GlOCPX structure also revealed the Arg154-Glu245 interdomain salt bridge, which breaks upon OCP photoactivation and interaction with PBs ^17^, and is missing in some OCP2 homologs ^37^.

A well-defined density revealed that the GlOCPX-embedded carotenoid is echinenone (ECH), accommodated between the NTD and CTD with the help of the conserved tyrosine and tryptophan residues very similar to SynOCP1 (Fig. 2F,G). Yet, we noticed that the ECH-binding tunnel of GlOCPX is characterized by an unusually high number of Met residues (8 instead of 5 in SynOCP1), especially abundant around the ketolated β-ring of ECH in the CTD (5 instead of 2 in SynOCP1) (Fig. 2F,G). This can be relevant for the OCP activity as a reactive oxygen species quencher ^11,56,57^.

A completely novel element observed in the GlOCPX structure is a short α-helix in the NTD that uniquely plugs the carotenoid tunnel (Fig. 2E and Fig. 3A,B). For example, this region is represented by a rather long flexible loop in SynOCP1 (Fig. 3A) and in the isolated NTD domain thereof (Fig. 3B) ^13^. Of note, despite it adopts such distinct conformations, sequences of this specific region of GlOCPX (**AP**A**P**KD**YQL**) and SynOCP1 (**AP**I**P**AG**YQL**) are very similar, which would complicate prediction of the presence of the α-plug structure in other OCPs directly from sequence. Unlikely the α-plug is caused by GlOCPX crystallization, as it is not involved in crystal contact formation.

**Fig. 3.**
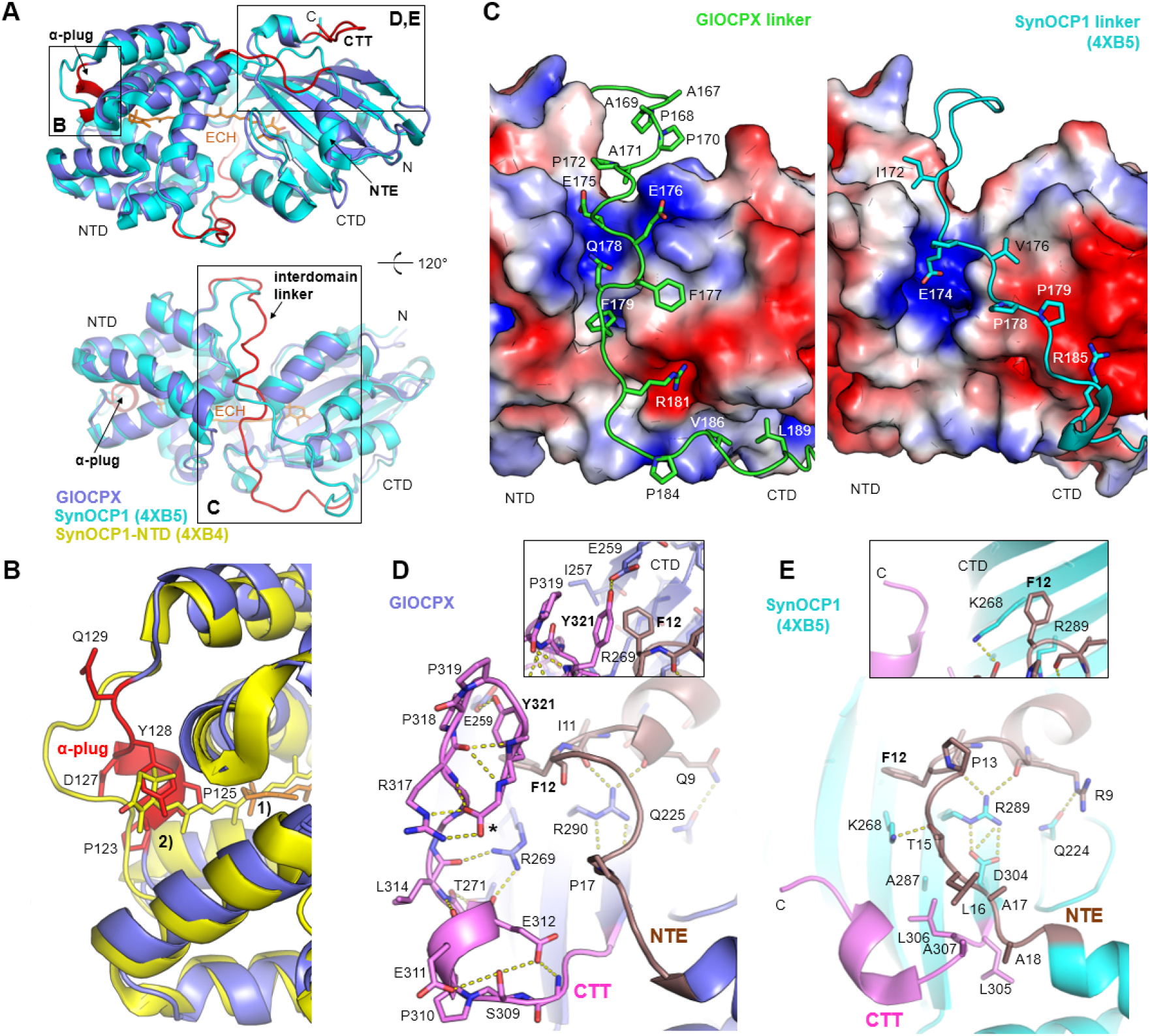
Distinct structural features of GlOCPX. A. Two opposite views of the GlOCPX structure (violet) overlaid with the SynOCP1 structure (cyan, PDB 4XB5) showing distinct structural features of GlOCPX (red). Echinenone (ECH) is shown by orange sticks. N-terminal (NTD) and C-terminal (CTD) domains, N-terminal extension (NTE), C-terminal tail (CTT), the interdomain linker and α-plug are labeled. Frames outline areas of special interest that are considered in panels B-E. B. Superposition of GlOCPX (violet cartoon, ECH - orange sticks) and the isolated CAN-bound NTD of SynOCP1 (yellow, PDB 4XB4) showing the shift of the carotenoid position from 1) to 2) and several residues of the α-plug (red, labeled) that restrict the tentative path for the carotenoid sliding in GlOCPX. C. Different interactions formed by the interdomain linkers of GlOCPX (left) and SynOCP1 (right). Key residues of the linkers are shown as sticks, the NTD and CTD are represented by surface electrostatics colored from red (−2 kT/e) to blue (+2 kT/e). D,E. Magnified views of the CTD region of GlOCPX (D) and SynOCP1 (E) revealing intricate interactions of the CTT (magenta) and NTE (chocolate) elements in GlOCPX securing its compact conformation. Asterisk denotes the C-terminal COOH group. Inserts show top views on the CTT/CTD interaction area and the involvement of GlOCPX-specific Tyr321 in CTT immobilization.

It is known that upon OCP1 photoactivation, carotenoid slides 12 Å deeper into the NTD from its original position (Fig. 3B), which determines spectral and functional properties of the signaling OCP^R^ state ^12,13,26^. However, given the location of the rather rigid α-plug in GlOCPX, which has a couple of Pro residues and a bulky Tyr128 side chain (Fig. 3B), the presumed carotenoid sliding would result in a severe clash. We therefore speculate that the α-plug shortens the carotenoid tunnel, which may have functional implications including a reduced photoproduct yield.

The GlOCPX structure is different from the SynOCP1 structure in the same areas that are incorrectly predicted by Alphafold2 (Fig. 2D and Fig. 3A). Besides the α-plug, the interdomain linker, which is known to actively participate in the OCP recovery ^15,38^, adopts a significantly different position in GlOCPX than in SynOCP1 (Fig. 3A,C). The GlOCPX linker features a remarkable pattern of several Pro residues interspaced by Ala residues (^167^APAPAP^172^), which albeit not forming direct contacts with the surface of GlOCPX, makes this linker subpart rather rigid (Fig. 3C). Another part of the GlOCPX linker makes tight contacts with the NTD and CTD based on complementary charges (e.g., Arg181, Glu175, Glu176 of the linker) or hydrophobic interactions (e.g., Phe177, Phe179, V186, L189 of the linker). In sharp contrast, interactions of the linker with the SynOCP1 domains are very scarce (e.g., changed interactions of Glu174 and Arg185 and hydrophobic interactions involving I172 and V176 of the linker). Overall, Fig. 3C shows that the interdomain linkers not only differ by the conformation, but also that the GlOCPX linker much tightly lies within the interdomain groove and stabilizes the OCP^O^. structure by multipoint interaction. This is in line with the more compact GlOCPX size observed on SEC (Fig. 1). Given the well-recognized role of the interdomain linker in OCP photocycle, our structure predicts that the GlOCPX photocycle is different from that of SynOCP1.

GlOCPX structure revealed a remarkable interplay of the NTE and CTT elements compared to the SynOCP1 structure, where almost no direct interactions between the NTE and CTT are found (Fig. 3D,E). First of all, the CTT of GlOCPX formed a peculiar, kinked loop structure maintained by two adjacent Pro residues in positions 318 and 319 and a range of intrachain interactions, including the salt bridge between the protein carboxyl group and the side chain of Arg137, as well as several H-bonds between the backbone O and N atoms (Fig. 3D). Furthermore, the apparently rigid CTT is fixed on the CTD via H-bonds to the CTD’s Arg269, Thr271 and Glu259, as well as a remarkable velcro-type contact of the Tyr321 side chain and Phe12 of the NTE. Phe12 makes a hydrophobic contact with the Cβ-Cγ-Cδ part of Arg269, whereby a triple interaction between the CTT, NTE and CTD is achieved (Fig. 3D). In striking contrast, most of these interactions are missing in SynOCP1, which has a shorter and apparently rather flexible CTT (Fig. 3E). While the NTE immobilization on the CTD appears similar in both proteins, the CTT of GlOCPX displaces the loop connecting the α1-helix of the NTE with the NTD, which is facilitated by a kink at Pro17 (Fig. 3D). Obviously, such an intricate network of chemical interactions involving the CTT and NTE of GlOCPX could not be predicted.

### GlOCPX photoactivity

The absorbance spectrum of GlOCPX reversibly transforms in response to blue LED illumination (Fig. 4A). While this is similar to SynOCP1 qualitatively, the relative photoactivity of GlOCPX is lower than SynOCP1 (Fig. 4B) and the kinetics of GlOCPX recovery after photoactivation is remarkably fast, much faster than SynOCP1 at a range of temperatures (Fig. 4C). For example, when at 5 °C full recovery took ∼30-35 min for GlOCPX, SynOCP1 recovered by less than 5% (Fig. 4C). However, at 35 °C, the recovery kinetics became comparable, and full recovery of either protein took <5 min (Fig. 4C). The temperature dependences of the recovery rate (ln*k*_R-O_), the so-called Arrhenius plots, revealed two linear segments with an inflection point at 20 °C for each OCP type. Such an inflection has been reported for OCP ^24,38^, and likely indicates that H-bonding and hydrophobic interactions, known to oppositely depend on temperature, contribute differently into the efficiency of restoration of the OCP^O^. state at temperatures lower or higher than 20 °C. Nevertheless, for both such segments on the Arrhenius plot, the R-O transition for GlOCPX is much less temperature-dependent (i.e, lower activation energy) compared to SynOCP1 (Fig. 4D). These observations are rationalized by the structural differences, in particular the more rigid and tightly bound interdomain linker and the peculiar NTE/CTT lock in the case of GlOCPX (Fig. 3).

**Fig. 4.**
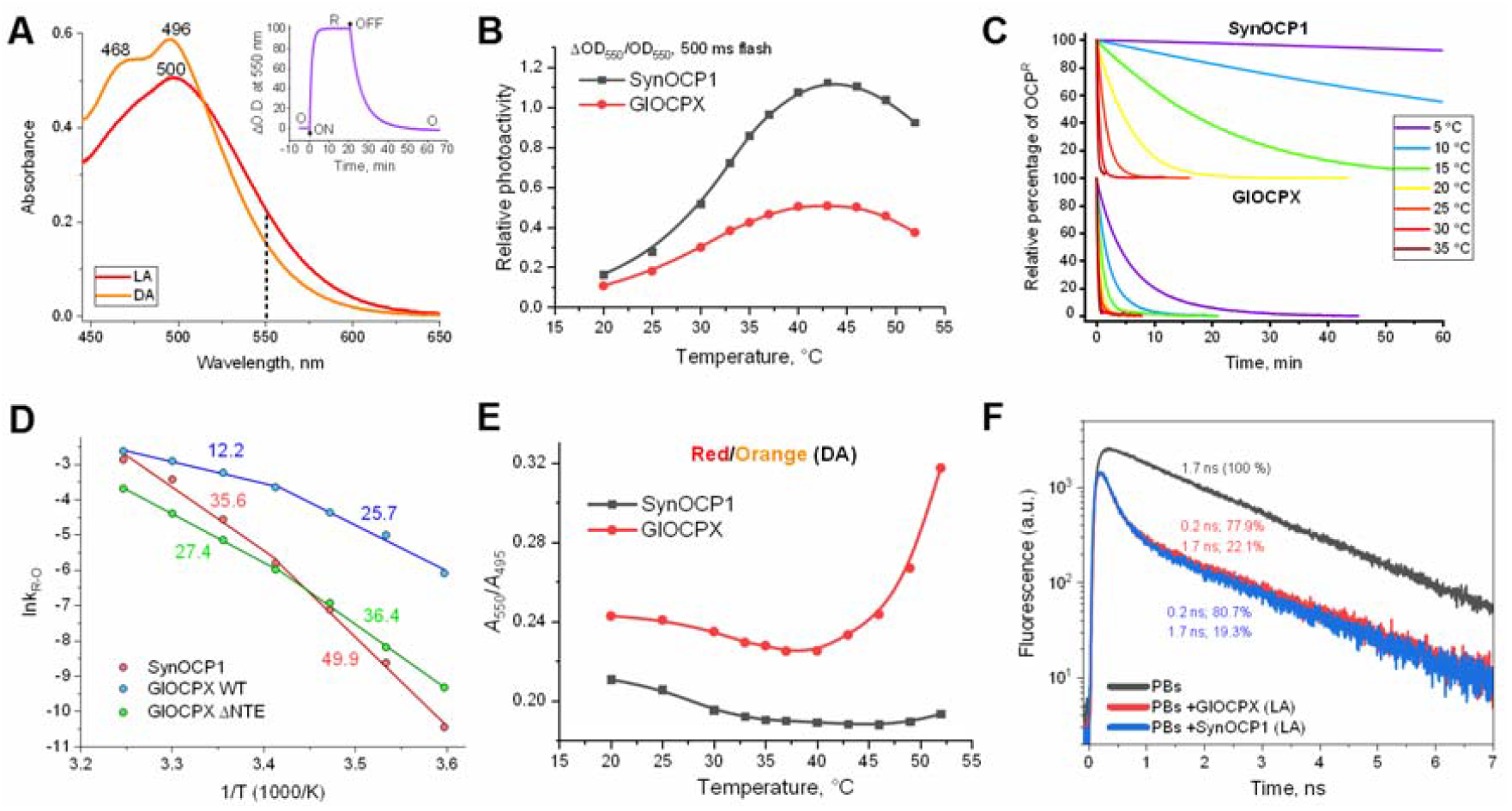
GlOCPX photoactivity studied by spectroscopy. A. Spectral changes of GlOCPX upon photoactivation by the blue LED. LA, light-adapted; DA, dark-adapted spectrum. A dashed line indicates where the photocycle is measured. Insert shows exemplary kinetics of O-R and R-O transitions for GlOCPX WT at 5 °C followed by changes of absorbance at 550 nm. Arrows indicate when the blue LED was switched on and off. B. Dependence of the relative photoactivity of SynOCP1 and GlOCPX on temperature. C. R-O transition of SynOCP1 and GlOCPX at different temperatures (color coded according to the legend). D. Arrhenius plots for GlOCPX WT and its ΔNTE mutant, compared with SynOCP1. Activation energies (*E*_A_) are indicated in kcal/mol as calculated for low-temperature and high-temperature segments independently. E. Influence of temperature on the yield of the red state in the compact dark-adapted state of GlOCPX or SynOCP1. F. Quenching of fluorescence of *Synechocystis* phycobilisomes upon the photoactivation of OCPs by 5 min light adaptation (LA) to blue LED. Fluorescence decay kinetics were recorded at 680 nm with excitation at 620 nm. The numbers indicate the characteristic lifetimes of the excited states and their contributions. Measurements were performed in 0.8 M phosphate buffer at 25 °C. The OCP/PBs concentration ratio was ∼40/1. All samples uniformly contained ECH as a cofactor.

Of note, the NTE deletion decelerated GlOCPX recovery and made its rate much more temperature-dependent (*E*a increased for both segments on the Arrhenius plot, Fig. 4D), while the inflection became less pronounced. This agrees well with the widely recognized role of NTE in stabilizing the compact OCP^O^. conformation ^15,16,55^ and indirectly indicates that the presence of the two segments on the Arrhenius plots of OCP is determined by a combinatorial effect of the interdomain linker and NTE in the activation energy of the R-O transition. The absence of the NTE obviously makes the OCP^O^. state restoration a challenge due to the lack of many stabilizing contacts in ΔNTE-GlOCPX (Fig. 3D). It simultaneously minimizes the inflection of the Arrhenius plot, revealing the predominant effect of the interdomain linker. Noteworthily, GlOCPX lacking NTE recovers to the compact state nearly as efficiently as SynOCP1 WT (Fig. 4D). It is therefore tempting to speculate that the combined effect of the interdomain linker and NTE in SynOCP1 is roughly balanced by the effect of the interdomain linker of GlOCPX alone and that these structural elements differently affect the temperature dependences of the R-O transition for any OCP. Remarkably, we found that the red-shift of the steady-state dark-adapted absorbance spectrum of GlOCPX relative to that of SynOCP1 (Fig. 1D) becomes more pronounced with increasing temperature and that this is reversible (Fig. 4E). Such changes suggest increased dynamics of the H-bonds between the carotenoid and protein in the CTD due to specific organization of the carotenoid-binding tunnel in GlOCPX (Fig. 2F).

Thus, the CTT/NTE lock and interdomain linker partly compensate for the less secure carotenoid immobilization in GlOCPX.

### GlOCPX quenches fluorescence of phycobilisomes from *Synechocystis*

Next, we estimated the efficacy of the interaction between GlOCPX and PBs from *Synechocystis. In vitro*, PBs require high concentrations of kosmotropes (e.g., 0.8 M phosphate) for maintaining PBs integrity and demonstrate the characteristic fluorescence lifetime of 1.7-1.8 ns. GlOCPX addition to PBs caused no changes in the intensity and lifetime of the PBs fluorescence without blue-LED illumination (Fig. 4F), indicating no interaction in the dark-adapted compact GlOCPX state, in which the PBs-binding sites are inaccessible. However, these PBs-binding sites may be equivalent for GlOCPX and SynOCP1, since upon the photoactivation time-resolved fluorimetry indicated an almost identical efficiency of PBs fluorescence quenching of both OCPs (substantial decrease of the 1.7 ns component contribution, see Fig. 4F).

### GlOCPX is not regulated by FRP from other cyanobacteria

SynOCP1 is regulated by its partner protein SynFRP, which efficiently accelerates R-O transition by binding to the photoactivated SynOCP1 state with the detached NTE and separated structural domains and by promoting the restoration of the compact OCP^O^. state ^28,30^. In SynOCP1, the main SynFRP binding site is located on the CTD and partly overlaps with the site of NTE attachment (SASBDB ID: SASDDG9, Fig. 5A) ^16,32^. Since the NTE removal reveals the FRP-binding site in SynOCP1, we created an analogous GlOCPX variant lacking its NTE (Fig. 5B), which could be purified as a photoactive protein. Surprisingly, we did not detect any significant effect of SynFRP on the photocycle of GlOCPX and ΔNTE-GlOCPX, indicating lack of functional interaction (Fig. 5C). However, the possibility of nonproductive SynFRP binding to GlOCPX remained, which required further testing of direct binding of FRP to the dark-adapted ΔNTE-GlOCPX.

**Fig. 5.**
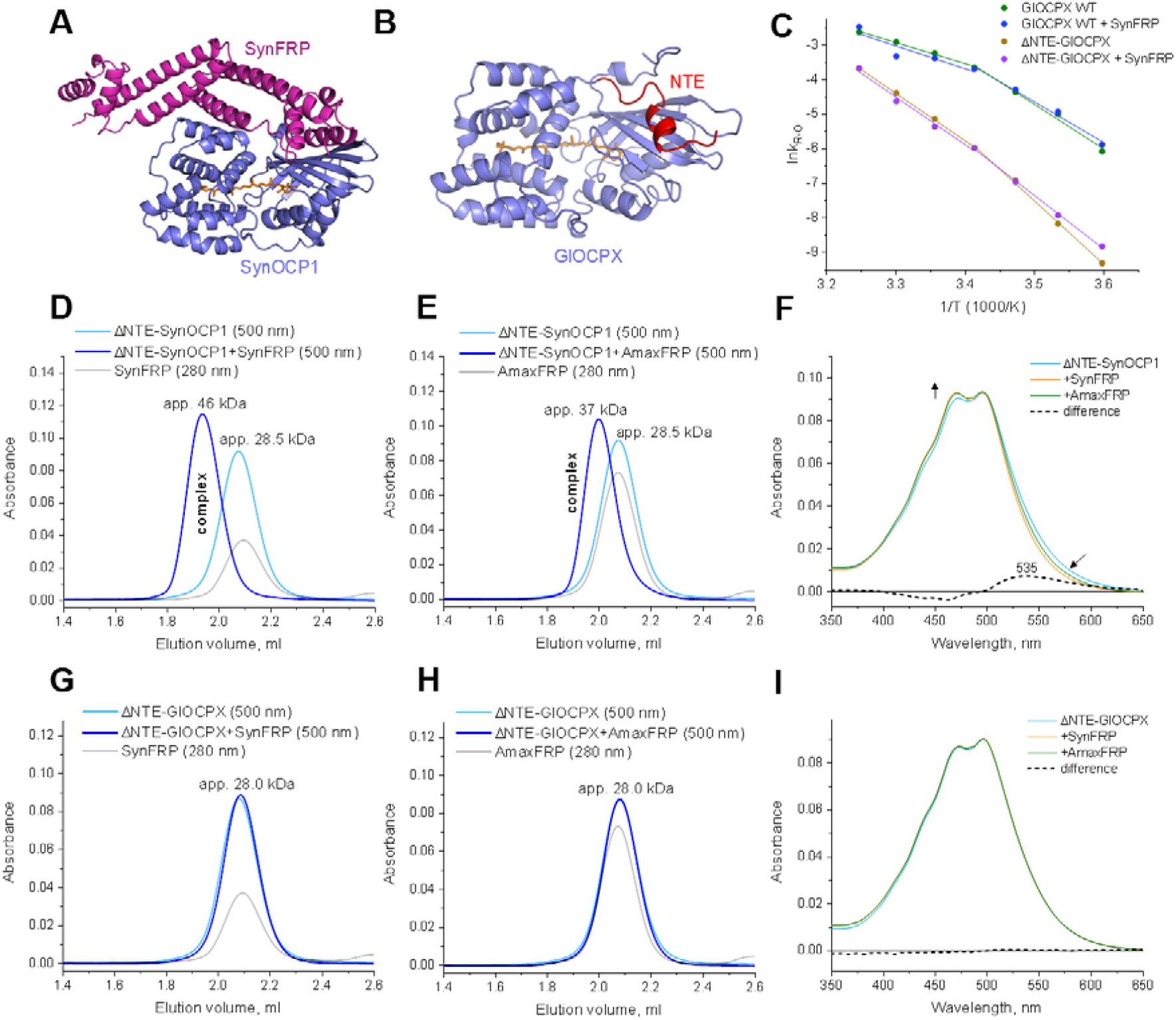
GlOCPX evades regulation by FRP from other cyanobacteria. A. Structural model of the SynFRP complex with ΔNTE-SynOCP1 (SASBDB ID SASDDG9 ^32^). B. Structural model of GlOCPX showing the NTE location (red). C. Arrhenius plots showing no effect of SynFRP on recovery kinetics for GlOCPX WT and its ΔNTE mutant. D-I. Physical interaction of either ΔNTE-SynOCP1 or ΔNTE-GlOCPX (20 μM each) with either SynFRP or AmaxFRP (40 μM each) studied by spectrochromatography using a Superdex 200 Increase 5/150 column and diode array detection. Effect of SynFRP or AmaxFRP on the absorbance spectrum (“oranging”) of ΔNTE-SynOCP1 (F) and ΔNTE-GlOCPX (I). Spectral changes associated with FRP binding are indicated by arrows. Dashed lines show difference spectra in the absence and in the presence of SynFRP.

As previously ^16,32^, mixing ΔNTE-SynOCP1 with SynFRP produced the stable complex eluting from SEC earlier than ΔNTE-SynOCP1 alone (Fig. 5D), which increased the apparent Mw from 28.5 kDa (ΔNTE-SynOCP1) to 46 kDa for the complex. Likewise, although causing a less pronounced shift on the elution profile, addition of FRP homolog from *Arthrospira* (*Limnospira*) *maxima* (AmaxFRP) also led to the ΔNTE-SynOCP1/FRP complexation (Fig. 5E). Previous work assigned the decreased apparent Mw of this complex (37 kDa) to a distinct stoichiometry of the AmaxFRP/ΔNTE-SynOCP1 interaction ^34^. The OCP/FRP interaction was also evident from the so-called “oranging” ^16,29^ of the ΔNTE-SynOCP1 absorbance spectrum under the action of either SynFRP or AmaxFRP (Fig. 5F). This spectral change reflects the formation of the complex, in which FRP prevents ΔNTE-SynOCP1 from photoactivation-related changes by forcing domain reassembly ^16^.

On the contrary, under identical conditions, no complex formation was detected if SynFRP or AmaxFRP were added to ΔNTE-GlOCPX (Fig. 5G,H), and the absorbance spectrum of the latter remained unchanged (Fig. 5I). These data indicate that GlOCPX is not recognized nor regulated by FRP orthologs from other cyanobacteria.

### Phylogenetic analysis suggests subdivision of OCPX into subclades

Having characterized the tentatively primordial OCPX from *Gloeobacter kilaueensis*, we analyzed how well our data represent the OCPX clade by constructing a phylogenetic tree rooted using sequences of OCP1 proteins with the resolved three-dimensional structure (Fig. 6A and Table 1). According to this analysis, the heterogeneous OCPX clade contains several subgroups where *Gloeobacteria* OCPX form a separate subgroup, the most distant from the OCP1 branch. Based on the observed heterogeneity, we suggest renaming of the OCPX clade into OCP3 (by analogy with OCP1 and OCP2), with its further subdivision into at least three subgroups. Given their structural peculiarities and origin, *Gloeobacteria* OCPX homologs are proposed to constitute a basal subclade, OCP3a, whereas its closest subclade is proposed to be named OCP3b (Fig. 6A). The rest OCP3 representatives are closest to OCP1 on the tree and are still a heterogeneous subgroup, which suggests their temporary naming as OCP3c (with a possibility of further subdivision into OCP3c, OCP3d, etc. in the future).

**Fig. 6.**
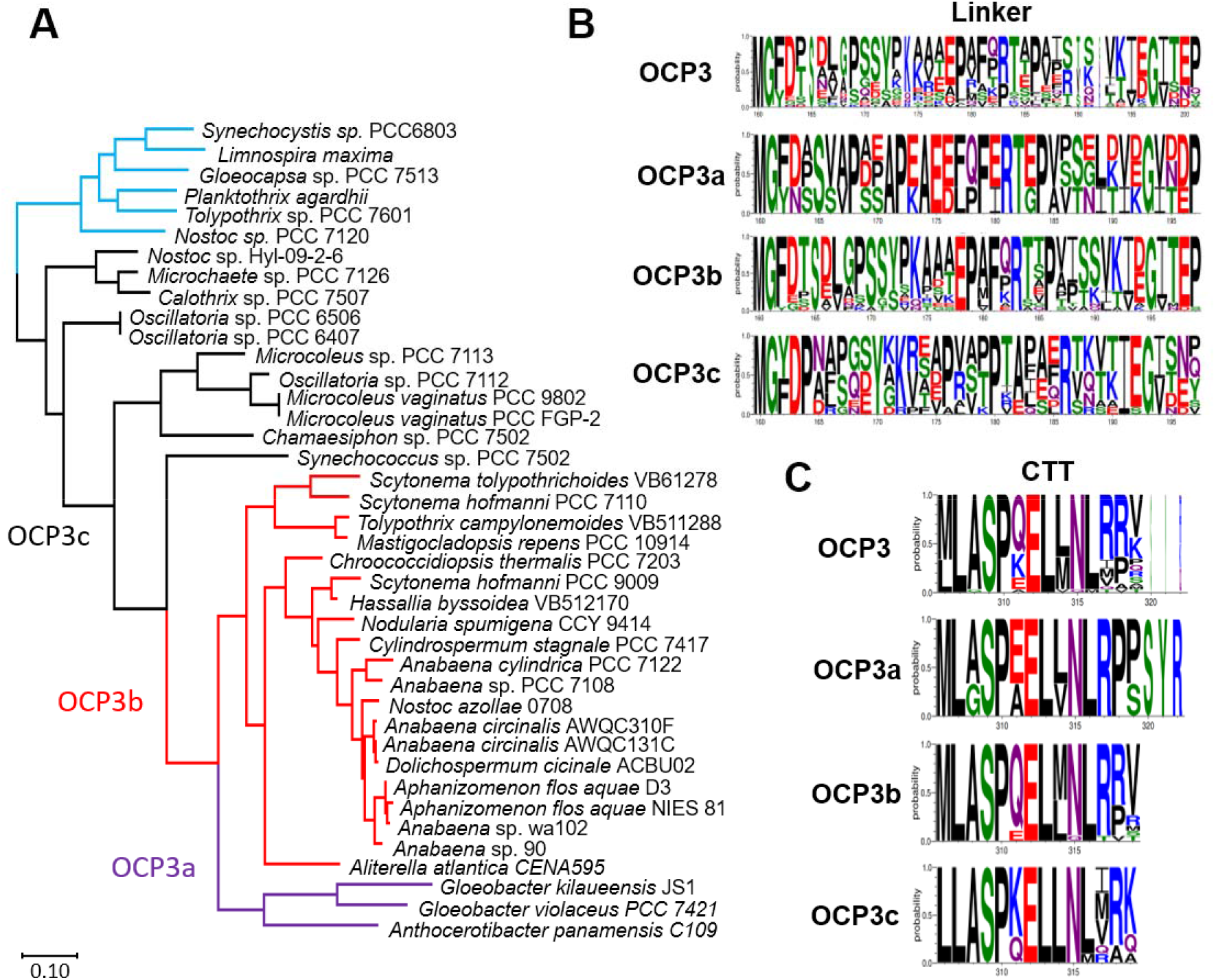
Phylogenetic analysis suggests subdivision of the heterogeneous OCPX clade. A. Phylogenetic tree of 36 single-copy OCPX and 6 OCP1 sequences with the known crystal structures (see also Table 1). The evolutionary history was inferred by using the Maximum Likelihood method based on the JTT matrix-based model ^68^. The tree with the highest log likelihood is shown. The tree is drawn to scale, with branch lengths measured in the number of substitutions per site. All positions containing gaps and missing data were eliminated. For the tree rooting, OCP1 representatives were used as an outgroup (coloured blue). B. WebLogo of linker sequences. C. WebLogo of CTT sequences. The default color coding is used.

The proposed subdivision enables a focused comparison of the key functionally relevant segments, fundamentally different in SynOCP1 and GlOCPX, i.e. the interdomain linker (Fig. 6B) and CTT (Fig. 6C). Apparently, the OCP3 subclades feature much clearer and informative logo diagrams for the corresponding sequences, compared to the blurred logo for the parent OCP3 (formerly, OCPX) clade combined. In particular, we found the distinctive distribution of the charged and hydrophobic residues in the linkers of the proposed subclades (Fig. 6B). The most similar are linker sequences of the OCP3a and OCP3b subclades. For instance, the segment **E**FQ**F**E**R**, that forms multisite interactions with the interdomain interface in GlOCPX structure (Fig. 3C), is representative for the OCP3a group and is very similar to the OCP3b linker consensus, **E**PA**F**Q**R** (Fig. 6B). The only OCPX proteins characterized earlier ^38^, from *Scytonema hofmanni* PCC 7110 and *Synechocystis* sp. PCC 7509, belong to the OCP3b subclade. Concerning the distribution of the charged and hydrophobic residues, the OCP3c linker consensus, having a Pro-rich central segment **K**…**P**(**V**/R).(**P**/T)**P**, is completely different from OCP3b and OCP3a consensus sequences but resembles the analogous OCP1 segment **R**V.E**PV**V(**P**/A)**P**. These observations closely follow the relationships determined by the phylogenetic tree. The similarity of the functionally relevant linkers of the OCP1 and OCP3c phylogenetic groups suggests that the latter contains homologs which are the closest to the OCP1/OCP2 ancestor.

The CTT segments are rather similar in all OCP3 sequences (Fig. 6C). However, the profound feature of the OCP3a CTT is its longer length and the presence of a Pro-Pro tandem, which in our structure forms the kink within a hairpin-like element, additionally stabilized by several interactions *in cis* (Fig. 3D). The unique antepenultimate Tyr residue forming the special lock with the NTE and CTD (Fig. 3D) is found only in the OCP3a subclade.

## Discussion

Recent bioinformatic analysis of the available OCP sequences suggested at least three families: OCP1, OCP2 and OCPX ^37^. OCPX has been proposed to represent the most diverged from OCP1 group. Evolutionary reconstruction based on the rooted phylogenetic tree of OCP sequences suggests at least two duplication events, giving initially OCPX and OCP1/OCP2 clades, with the latter splitting off at a later stage ^38^. In the aftermath, only OCP1 acquired the regulation by FRP ^37^, which accelerates R-O conversion and can be independently regulated by varying expression levels ^28^. It is unclear whether the OCP1/FRP system confers any adaptational advantages compared to OCP2 and OCPX, however, the majority of cyanobacteria encode a single OCP1 copy and an adjacent FRP gene.

Of note, neither OCP1 nor FRP are found in *Gloeobacteria*, which is a sister group relative to all other cyanobacteria and which contains exclusively OCPX ^37^. One can assume that some *Gloeobacter* branched off the other cyanobacteria long before the duplication and subfunctionalization of OCP yielding extant OCP1, OCP2 and OCPX groups, and thereby retained a predominant number of ancestral characteristics. From this perspective, *Gloeobacter* OCPX is the best possible extant version of an ancestor OCP. Importantly, *Gloeobacter* is capable of blue-light-triggered NPQ and echinenone biosynthesis and GlOCPX was evidenced at transcript and protein levels ^45,49^, which strongly indicates that it is a protein functional *in vivo*.

Our detailed comparison of the canonical SynOCP1 and its most distant counterpart OCPX from *Gloeobacter kilaueensis* JS1 (50.5% identity) revealed interrelated structural and functional differences. The O-R and R-O transitions for GlOCPX are less temperature-dependent and have lower activation energies, which may be relevant for extremophilic OCPX-containing cyanobacteria (e.g., *Chroococcidiopsis thermalis*). Peculiarities of the linker and CTT revealed by the GlOCPX crystal structure can explain a considerable decrease of *E*a of the R-O transition. This is in line with the faster recovery kinetics (compared to that of OCP1 and OCP2) of the two OCPX variants described earlier ^38^ and is apparently associated with the commonality of their linker sequences [^176^**E**FQ**FERT**E**P**^184^ (GlOCPX) or ^176^**E**P.**F**(**E/**P)**R**(D**/T**).**P**^184^ (SynOCPX and ScyOCPX)] (Supplementary Fig. 3A). One of the most controversial points remains the oligomeric state of OCP. SynOCP1 has been described either as a monomer ^29,32^ or a dimer ^37^. Notwithstanding, all six OCP1 homologs crystallized so far formed similar crystallographic dimers (Table 1). For SynOCPX and ScyOCPX, a significant tendency for dimerization *in vitro* has been reported ^38^. In contrast, GlOCPX shows a minor dimeric fraction even at ∼20 mg/ml concentration and is mostly represented by stable monomers in solution and in crystals. Of note, in contrast to GlOCPX having the Glu25 residue apparently incompatible with the dimerization observed in SynOCP1 structures (Table 1), both SynOCPX and ScyOCPX contain dimerization-compatible alanines in the corresponding position, just like SynOCP1 (Supplementary Fig. 2 and 3).

The inability of GlOCPX to interact physically and functionally with FRP from other cyanobacteria shown here is in line with the evolutionary considerations that the OCP-FRP interaction emerged relatively recently, and only for OCP1 homologs ^37,38^. This marked difference is likely accounted for by the scattered amino acid replacements acquired during the OCP1 evolution. At the same time, surprisingly, GlOCPX is as efficient quencher of fluorescence of PBs from *Synechocystis* as SynOCP1. In the absence of FRP in *Gloeobacteria*, the recovery of photosynthetic activity after GlOCPX photoactivation would obviously require considerable time unless alternative mechanisms are involved. As we showed earlier ^58^, CTDH is capable of performing a function similar in effect to FRP by temporarily arresting the ketocarotenoid via protein-to-protein carotenoid transfer. Given that at least three sequenced *Gloeobacter* species contain a CTDH homolog (Fig. 1B), this possibility warrants further investigation.

The ability of ScyOCPX ^38^ and GlOCPX to quench *Synechocystis* PBs indicates that an OCP-dependent NPQ mechanism emerged in cyanobacteria long before *Gloeobacteria* branched off from the crown cyanobacteria around 2 Ga years ago at the dawn of the oxygenic photosynthesis. Interestingly, GlOCPX and SynOCP1 share the semiconserved interdomain salt bridge involving Arg155 that was proposed to play a role in photoactivation and PBs quenching ^17^.

Analysis of the OCPX clade is complicated by its high heterogeneity ^37^. The current model of the OCP origin implies that before the fusion, the OCP domains represented individual proteins whose heterocomplex became photoactive ^37^. In the light of this model, the linker of the full-size OCP is the most recently acquired element ^37^. While we identified the highly specialized linker in GlOCPX, the linker sequence appears too variable if OCPX homologs are considered collectively (Fig. 6B, top). A similar, albeit less pronounced, heterogeneity is revealed by the C-terminal tail consensus for OCPX homologs (Fig. 6C).

To detect regularities in the OCPX organization, we revisited the earlier phylogenetic analysis of OCPX homologs ^37,38^. First of all, we discarded OCPX homologs co-existing with OCP1 or OCP2 in cyanobacterial genomes because their independent functionality a priori is unclear. Fortunately, while different combinations of OCP homologs may indeed be present, the majority of cyanobacterial genomes encode only a single OCP copy ^37^. It is likely that only in a few cases a genuine subfunctionalization of OCP homologs took place, as exemplified by *Tolypothrix* sp. PCC 7601, which contains OCP1 and OCP2 that are differently expressed and regulated ^37^. In a majority of cyanobacterial genomes, redundant OCP copies are thought to have been eliminated ^37^. Nonetheless, OCPX-containing cyanobacteria unite species where i) an OCP1/OCP2 ancestor has never existed, ii) has been eliminated, or iii) an OCP1 or OCP2 has been eliminated after the emergence. These scenarios likely made the heterogeneity of the OCPX group more profound.

Based on structural, functional and phylogenetic analysis, we proposed renaming of the heterogeneous OCPX clade into OCP3 and its subdivision into subclades, OCP3a, OCP3b and OCP3c (Fig. 6A), reflecting the most distant from OCP1 position of the *Gloeobacteria* OCP3a subclade and distinctive differences in the functionally relevant interdomain linkers. While two OCP3b ^38^ and one OCP3a (this work) representatives are described, the OCP3c subclade still remains uncharacterized and less homogeneous. According to our phylogenetic analysis and sequences of their functional elements, OCP3c homologs represent an intermediate evolutionary step between OCP3 and an OCP1 ancestor (Fig. 6A). Further investigation could help better understand the origin of OCP1 homologs, which became successful and considerably displaced other OCP variants.

## Materials and methods

### Proteins

cDNA corresponding to GlOCPX residues M1-R322 (Uniprot U5QHX0) was codon-optimized for expression in *E. coli*, synthesized by Integrated DNA Technologies (Coralville, Iowa, USA) and cloned into the pET28b-His-3C vector (kanamycin resistance) using the *NdeI* and *XhoI* restriction sites. The selected design left extra N-terminal GPH… residues after cleavage by specific rhinovirus 3C protease.

GlOCPX mutants E25A, E25C and ΔNTE (removal of the first M1-P17 residues) were obtained by the megaprimer method using Q5 (NEB) polymerase, corresponding mutagenic primers E25A-rev (5’-ATTAAACTCCGCTAAAATAGC-3’), E25C-rev (5’-ATTAAACTCGCATAAAATAGC-3’), ΔNTE-for (5’-ATTACATATGAGTCGCATCCCAGC-3’) and the wild-type plasmid as template. SynFRP (Uniprot P74103), AmaxFRP [Uniprot B5W3T4 (*Limnospira* (formerly, *Arthrospira*) *maxima* CS-328), coincides with Uniprot H1W9V5 (*Limnospira indica*) and K1X0E1 (*Arthrospira platensis* C1)], SynOCP1 (Uniprot P74102) and its ΔNTE mutant (lacking residues M1-F12 from the N-terminus) were obtained in previous work ^32,34^. All constructs were verified by DNA sequencing in Evrogen (Moscow, Russia) and transformed into C41(DE3) *E*.*coli* cells already carrying the pACCAR25ΔcrtXZcrtO plasmid (chloramphenicol resistance), which harbors the gene cluster including crtY, crtI, crtB, crtE and crtO sequences from *Erwinia uredovora* for ECH expression ^25^. Alternatively, for making GlOCPX(CAN) protein, co-expression with the β-carotene ketolase crtW (ampicillin resistance) on the background of the pACCAR16ΔcrtX plasmid (synthesizing β-carotene with the gene cluster crtY, crtI, crtB, crtE; chloramphenicol resistance) was used ^10,25^. IPTG-induced protein expression lasted overnight at 30 °C. In the case of GlOCPX(CAN) expression 0.01% arabinose was added to culture for induction of CrtW (CAN production). Proteins were purified according to the unified scheme consisting of subtractive immobilized metal-affinity and size-exclusion chromatography to electrophoretic homogeneity assessed by SDS-PAGE. Protein fractions were pooled together according to their Vis/UV absorbance ratios, 1.9 for GlOCPX(CAN) and 1.7 for GlOCPX(ECN). Purified proteins were stored frozen at -80 °C.

Disulfide trapping of the GlOCPX-E25C mutant dimer or monomer (concentrated SEC fractions) was achieved by dialyzing samples at ∼3 mg/ml protein concentration against 50 mM Tris-HCl buffer (pH 7.6) with GSH/GSSG (1 mM each) at 4 °C for 50 days. At different time points the extent of oxidation was assessed by SDS-PAGE in the absence or the presence of βME.

Phycobilisomes were isolated from *Synechocystis* cells as described earlier ^59^.

### Spectrochromatography

SEC with diode-array detection was used to analyze OCP species and the possibility of OCP/FRP interaction. Samples (50 μl) were loaded on a Superdex 200 Increase column (GE Healthcare, Chicago, Illinois, USA) pre-equilibrated with a 20 mM Tris-HCl buffer, pH 7.6, containing 150 mM NaCl and run by a Varian ProStar 335 HPLC system (Varian Inc., Melbourne, Australia). During the runs, absorbance in the 240-900 nm range was recorded with 1-nm precision (4 nm slit width) and a 5 Hz frequency. Alternatively, spectrochromatography was used during the final preparative SEC run on a Superdex 75 column (GE Healthcare). Column size is specified in figure legends. Diode-array data were converted into .csv files using a custom-built Python script and processed into contour plots using Origin 9.0 (Originlab, Northampton, MA, USA). Each experiment was repeated at least three times, and the most typical results are presented.

### SEC-MALS

Size-exclusion chromatography coupled to multi-angle light scattering (SEC-MALS) of GlOCPX or SynOCP1 samples (5 mg/ml in 50 μl) was performed on a Superdex 200 Increase 10/300 column (GE Healthcare) connected to a Prostar 335 detector (Varian, Australia) and a miniDAWN detector (Wyatt Technology, USA). The column was equilibrated with filtered (0.1 μm) and degassed 20 mM Tris-HCl buffer, pH 7.6, containing 150 mM NaCl. Flow rate was 0.8 ml/min. Data were analyzed in ASTRA 8.0 (Wyatt Technology, USA) using dn/dc = 0.185 and protein extinction coefficients ε(0.1%) at 280 nm equal to 1.40 (GlOCPX(ECH)) and 1.25 (SynOCP1(ECH)). These extinction coefficients took into account that ECH absorbance at 280 nm is ∼⅛ of its absorbance at 500 nm. To account for small laser wavelength (660 nm) absorbance, the laser intensity was corrected by forward laser power, which did not exceed a 2.5% drop for OCP.

### Crystallization of GlOCPx

Initial crystallization screening of the GlOCPX-E25A mutant was performed using a robotic crystallization system (Oryx4, Douglas Instruments, UK) and commercially available crystallization screens (Hampton Research, USA) by sitting drop vapor diffusion at 15 °C. The protein concentration was 22.2 mg/ml in 20 mM Tris-HCl buffer, pH 7.6. The drop volume was 0.3 µl with a 1:1 and a 2:1 protein-to-precipitant ratio. Well-shaped crystals were obtained in 0.2 M Ammonium sulfate, 0.1 M MES monohydrate pH 6.5, 30% w/v Polyethylene glycol monomethyl ether 5000.

### Data collection, structure determination and refinement

GlOCPx E25A mutant crystals were briefly soaked in 100% paratone oil (Hampton Research, USA) immediately prior to diffraction data collection and flash-frozen in liquid nitrogen. The data were collected at 100K at ID30A-3 beamline (ESRF, France). Indexing, integration and scaling were done using Dials ^60^ (Table 1). The structure was solved by molecular replacement with MOLREP ^61^ using the predicted Alphafold2 ^53^ structure as an initial model. The refinement was carried out using BUSTER ^62^ and REFMAC5 ^63^. The isotropic individual atom B-factors as well as hydrogens in riding positions were used during the refinement. The visual inspection of electron density maps and manual model rebuilding were carried out in COOT ^64^.

### Small-angle X-ray scattering

GlOCPX WT (35 µl, 17.5 mg/ml) or its E25A mutant (40 µl, 13.5 mg/ml) were loaded onto a Superdex 200 Increase 3.2/300 column (Cytiva) and eluted at a 0.075 ml/min flow rate while the SAXS data (I(s) versus s, where s = 4πsinθ/λ, 2θ is the scattering angle and λ=0.96787 Å) were measured at the BM29 beam line (ESRF, Grenoble, France) using a Pilatus 2M detector (data collection rate 0.5 frame/s; experiment session data DOI 10.15151/ESRF-ES-642726753). The buffer included 20 mM Tris-HCl, pH 7.6, and 150 mM NaCl. SAXS frames acquired during the SEC profile were processed in CHROMIXS ^65^ and the resulting SAXS profiles were used for validation using CRYSOL ^66^ and models of the GlOCPX E25A monomer and the tentative dimer built by structural overlay with the crystallographic SynOCP1 dimer (PDB 4XB5) ^13^.

### OCP photocycle followed by absorbance spectroscopy

To study the photocycle of different OCP forms, OCP samples (10 μM) in the absence or the presence of SynFRP (20 μM) were transferred to a 1 cm quartz cuvette and the steady-state absorbance spectrum was continuously registered. For photoconversion, a blue light-emitting diode (M455L3, Thorlabs, USA) with a maximum emission at 445 nm was used. The temperature of the sample was stabilized by a Peltier-controlled cuvette holder Qpod 2e (Quantum Northwest, USA). Each experiment was repeated at least three times, and the most typical results are presented.

### Time-resolved fluorescence measurements

PBs fluorescence decay kinetics with picosecond time resolution were recorded with a time- and wavelength-correlated single photon counting setup based on a HMP-100-40 detector and a SPC-150 module (Becker&Hickl, Germany). Fluorescence was excited at 620 nm by 150 fs flashes generated by optical parametric generator TOPOL (Avesta Project Ltd., Russia) PBs emission was recorded at 680 nm. Fluorescence decay was approximated by a sum of two exponential decay functions with the SPCImage (Becker&Hickl, Germany) software package, considering the incomplete decay of the states with long lifetimes at 80 MHz repetition rate. The temperature of the sample was controlled by a Qpod 2e (Quantum Northwest, USA) cuvette holder. Photoactivation of OCP was triggered by a blue LED (445 nm, 200 mW, Thorlabs, USA).

### Phylogenetic and bioinformatic analysis

For simplicity, we chose OCPX from cyanobacterial genomes where only a single copy of such a gene is present. Protein sequences of 34 OCPX homologs were retrieved from the dataset available in ^37^, with a lower length cut-off of 318 residues, which is apparently the minimal full length of OCPX. For the outgroup we additionally picked six OCP1 representatives with the known crystal structure (Table 1). This set of 34 OCPX and 6 OCP1 was aligned using T-COFFEE ^67^. A tree was built using MEGAX with the Maximum Likelihood method based on the JTT matrix-based model ^68^. The analysis involved 40 amino acid sequences. All positions containing gaps were eliminated. There were a total of 306 positions in the final dataset. Initial tree(s) for the heuristic search were obtained automatically by applying Neighbor-Join and BioNJ algorithms to a matrix of pairwise distances estimated using a JTT model, and then selecting the topology with a superior log likelihood value. The tree was rooted using OCP1 sequences as an outgroup. Family clades were determined by manual inspection.

## Abbreviations used

CAN: canthaxanthin
ECH: echinenone
hECH: 3-hydroxyechinenone
CTD: C-terminal domain
CTDH: C-terminal domain homolog
DA: dark-adapted
FRP: fluorescence recovery protein
OCP: Orange Carotenoid Protein
GlOCPX: *Gloeobacter kilaueensis* OCP clade X
HCP: helical carotenoid protein
IMAC: immobilized metal-affinity chromatography
IPTG: isopropyl-β-thiogalactoside
LA: light-adapted
LED: light-emitting diode
NPQ: nonphotochemical quenching
NTD: N-terminal domain
NTE: N-terminal extension
PBs: phycobilisomes
SAXS: small-angle X-ray scattering
SDS-PAGE: sodium dodecyl sulfate-polyacrylamide gel electrophoresis
SEC: size-exclusion chromatography
SEC-MALS: size-exclusion chromatography coupled to multi-angle light scattering
SynOCP1: *Synechocystis* sp. PCC 6803 OCP clade 1.

## Data availability

The refined model and structure factors have been deposited in the Protein Data Bank under the accession code 8A0H. All materials are available from the corresponding author upon reasonable request.

## Acknowledgements

The authors are grateful to Prof. Kai-Hong Zhao (Huazhong Agricultural University, Wuhan, China) for the pBAD-crtW plasmid, to Ilia Chetviorkin for the Python script for processing the diode-array data, to Dr. Dmitrii Zlenko for providing phycobilisome preparations and to Anton Popov and Dmitry Soloviov for help with SAXS data collection at the BM29 beamline (ESRF, Grenoble, France; experiment session data DOI 10.15151/ESRF-ES-642726753). The study was supported by the Ministry of Science and Higher education of the Russian Federation in the framework of the Agreement no. 075-15-2021-1354 (07.10.2021). Carotenoprotein expression and purification was supported by the Russian Foundation for Basic Research grant (no. 20-54-12018). Spectroscopic characterization of OCP-PBs interactions was supported by the Russian Science Foundation (grant number 21-44-00005).

## Author contributions

NNS – designed experiments, initiated and supervised the study; YBS, AOZ, NNS – expressed and purified proteins; LAV - crystallized proteins; YBS, AOZ, EGM, NNS – performed experiments; KMB, NNS - collected X-ray data and solved crystal structure; NNS - processed and analyzed SAXS data; YBS, KMB, EGM, NNS – analyzed data and discussed the results; NNS wrote the paper with input from YBS.

## Supplementary information

**Supplementary Fig. 1.**
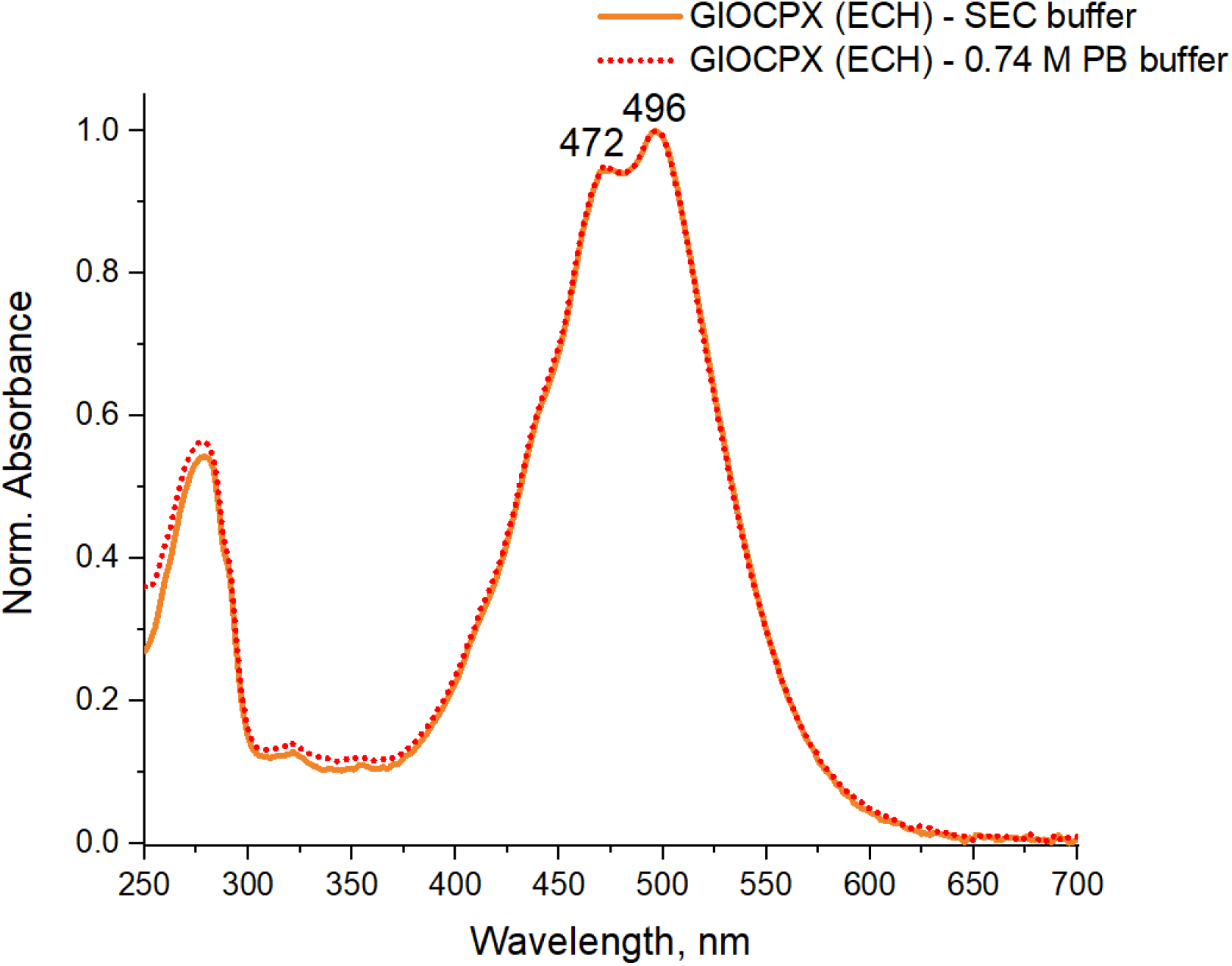
Effect of phosphate on the absorbance spectrum of GlOCPX.

**Supplementary Fig. 2.**
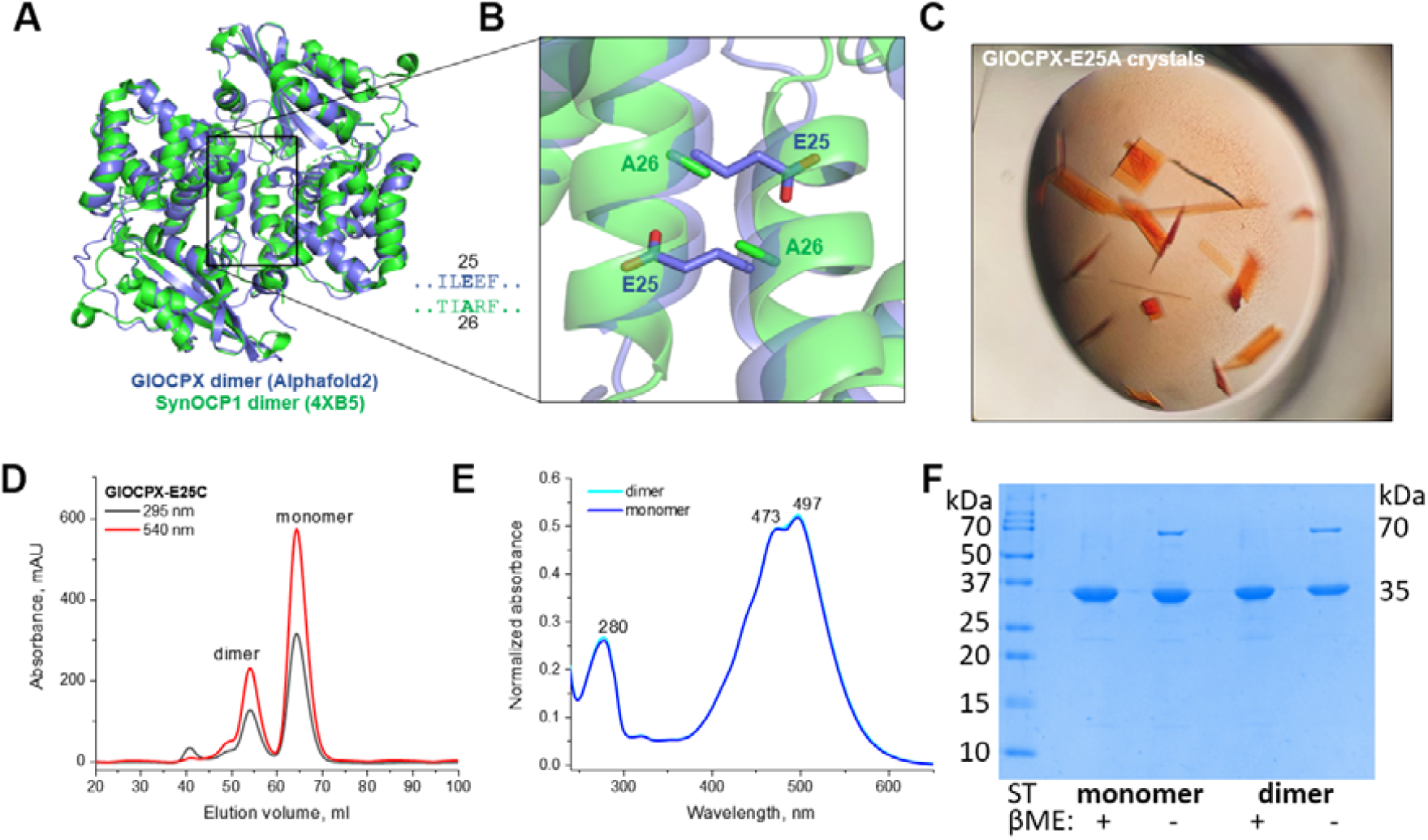
Engineering of the tentative dimer interface affects GlOCPX dimerization and crystallization. A. Crystallographic SynOCP1 dimer (PDB 4XB5) and an Alphafold2 model of GlOCPX were used to build a tentative model of the GlOCPX dimer. B. A magnified view of the interface showing the Ala26 residues of SynOCP1 and a severe clash caused by the Glu25 residues of GlOCPX in the equivalent position. C. Only the GlOCPX-E25A mutant and not the wild-type GlOCPX crystallized. D. Preparative SEC of the GlOCPX-E25C mutant on a Superdex 75 16/600 column showing two major peaks of the dimer and monomer. The normalized absorbance spectra of these two species are indistinguishable (E). F. Disulfide trapping of the GlOCPX-E25C dimer starting from either monomeric or dimer fraction (50-day dialysis under non-reducing conditions at 4 °C), analyzed by SDS-PAGE in the absence (-) or the presence (+) of β-mercaptoethanol (βME).

**Supplementary Fig. 3.**
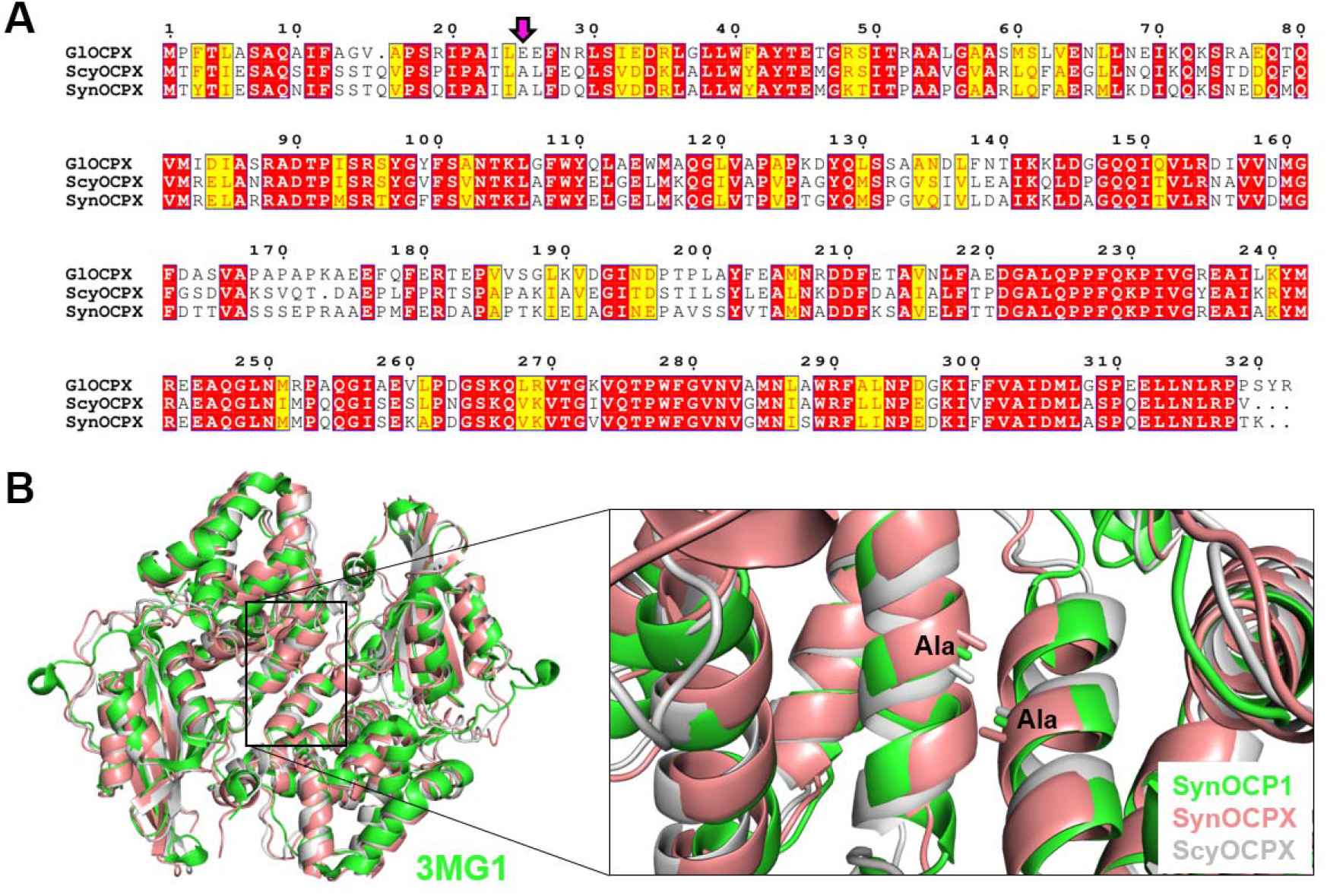
Alignment of sequences and structural models for selected OCP1 and OCPX homologs. A. Alignment of amino acid sequences of OCPX from *Gloeobacter kilaueensis* JS1 (IMG ID 2558546973) analyzed in this work and OCPX from *Scytonema hofmanni* sp. PCC 7110 OCPX (IMG ID 2551963320) and *Synechocystis* sp. PCC 7509 OCPX (IMG ID 2517697059) analyzed earlier ^38^. Sequence identifiers are indicated according to the Integrated Microbial Genomes and Microbiomes (IMG) database (https://img.jgi.doe.gov). A magenta arrow indicates the key position in the tentative dimer interface that is different in GlOCPX and the two other OCPX homologs. Red highlights the identical, whereas yellow highlights similar residues. B. Crystallographic SynOCP1 dimer (PDB ID 3MG1) overlaid with the structural models of OCPX from *Scytonema hofmanni* sp. PCC 7110 (ScyOCPX) and *Synechocystis* sp. PCC 7509 (SynOCPX) as predicted by Alphafold2. The magnified view on the right illustrates that like SynOCP1, both SynOCPX and ScyOCPX contain Ala in the dimer interface position, in which GlOCPX uniquely contains Glu25 residues incompatible with the dimer formation.

